# Evaluating the mouse neural precursor line, SN4741, as a suitable proxy for midbrain dopaminergic neurons

**DOI:** 10.1101/2023.01.23.525270

**Authors:** Rachel J. Boyd, Sarah A. McClymont, Nelson B. Barrientos, Paul W. Hook, William D. Law, Rebecca J. Rose, Eric L. Waite, Dimitrios Avramopoulos, Andrew S. McCallion

## Abstract

To overcome the ethical and technical limitations of *in vivo* human disease models, the broader scientific community frequently employs model organism-derived cell lines to investigate of disease mechanisms, pathways, and therapeutic strategies. Despite the widespread use of certain *in vitro* models, many still lack contemporary genomic analysis supporting their use as a proxy for the affected human cells and tissues. Consequently, it is imperative to determine how accurately and effectively any proposed biological surrogate may reflect the biological processes it is assumed to model. One such cellular surrogate of human disease is the established mouse neural precursor cell line, SN4741, which has been used to elucidate mechanisms of neurotoxicity in Parkinson disease for over 25 years. Here, we are using a combination of classic and contemporary genomic techniques – karyotyping, RT-qPCR, single cell RNA-seq, bulk RNA-seq, and ATAC-seq – to characterize the transcriptional landscape, chromatin landscape, and genomic architecture of this cell line, and evaluate its suitability as a proxy for midbrain dopaminergic neurons in the study of Parkinson disease. We find that SN4741 cells possess an unstable triploidy and consistently exhibits low expression of dopaminergic neuron markers across assays, even when the cell line is shifted to the non-permissive temperature that drives differentiation. The transcriptional signatures of SN4741 cells suggest that they are maintained in an undifferentiated state at the permissive temperature and differentiate into immature neurons at the non-permissive temperature; however, they may not be dopaminergic neuron precursors, as previously suggested. Additionally, the chromatin landscapes of SN4741 cells, in both the differentiated and undifferentiated states, are not concordant with the open chromatin profiles of *ex vivo*, mouse E15.5 forebrain- or midbrain-derived dopaminergic neurons. Overall, our data suggest that SN4741 cells may reflect early aspects of neuronal differentiation but are likely not a suitable a proxy for dopaminergic neurons as previously thought. The implications of this study extend broadly, illuminating the need for robust biological and genomic rationale underpinning the use of *in vitro* models of molecular processes.

## BACKGROUND

*in vitro* cellular surrogates present an excellent opportunity for elucidating the molecular mechanisms behind human disease without the ethical and technical limitations of *in vivo* systems. As such, most studies of human disease that employ genomic or cellular manipulations or assays that require high cell quantity and quality, are often conducted *in vitro* to ensure biological and statistical robustness[1–3]. For example, *in vitro* models are frequently employed in studies of the role of genomic regulation in human disease, identification of candidate genes and regulatory elements and evaluation of their functional characteristics through genetic manipulations and high-throughput assays[4–7]. As genome-wide association studies (GWASs)continue to indict human disease-associated variants, it is becoming evident that most of them lie within non-coding regions of the genome[8]. Such regions frequently represent cis-regulatory elements (CREs), required for the transcriptional modulation of cognate genes. The assays required to evaluate their function[4,9,10], or connect CREs with the promoters they modulate[11], often require large cell numbers, making, *in vitro* cellular systems the preferred strategy.

Prioritizing non-coding GWAS variants and disease-relevant sequences for extensive investigation requires knowledge of their chromatin accessibility status. Open chromatin is prone to harbor functional sequences; and since chromatin accessibility profiles vary across cell types and developmental time, it is important to prioritize disease-associated variants that lie within open chromatin regions in the disease-relevant cell type(s) [8,12,13]. It is also critical to functionally evaluate the biological consequences of disease-associated variation, test the efficacy of potential therapeutics, and observe the effects of disease-relevant insults in the appropriate cellular context[8,14,15]. Therefore, when studying disease associated variation, the most effective *in vitro* cellular surrogates should ideally mimic the chromatin architecture and transcriptional profiles of the *in vivo* cell types affected by disease.

In Parkinson disease (PD), midbrain (MB) dopaminergic (DA) neurons in the substantia nigra (SN) are the primary affected cell type[16]. Preferential degeneration of these neurons elicits a progressive neurodegenerative disorder characterized by motor deficits[16]. As the second most common neurodegenerative disorder, affecting approximately 1% of adults over 70 years old[17,18], PD is the focus of extensive research efforts. As such, various cell lines have been used as *in vitro* proxies of MB DA neurons to study the cellular impacts of PD-relevant insults, as well as candidate PD-associated sequences, their functions, and their potential as therapeutic targets[19].

One such cell line, SN4741, is reported to be a clonal DA neuronal progenitor line that was established in 1999 from mouse embryonic day 13.5 (E13.5) SN tissue[20]. The SN was dissected from transgenic mice containing 9.0 kb of the 5’ promoter region of tyrosine hydroxylase (*TH*), fused to the temperature-sensitive mutant Simian Virus 40 T antigen (SV40Tag-tsA58) oncogene[20]. The goal of this *TH* promoter transgene was to enable selective acquisition of DA neurons, while the purpose of the SV40Tag oncogene was to facilitate conditional immortalization of the cell line. The temperature sensitive mutant form of this immortalizing gene (tsA58) should permit uncontrolled differentiation and proliferation at the permissive temperature (33°C), maintain cells in an undifferentiated state at 37°C, and since tsA58 displays diminished activity at 39°C, it should direct differentiation that more closely resembles primary cells when the culture is shifted to this non-permissive temperature[20].

As an established mouse neural precursor line, SN4741 cells have since been used to elucidate mechanisms of neurotoxicity in PD[21–25], test the efficacy of therapeutic targets against PD relevant insults[26,27], and assay the impacts of PD-associated genetic mutation[28,29] and transcriptional regulation[30–32]. Important technological advances have also arisen since the genesis and implementation of the SN4741 cell line, including chromatin conformation capture technologies[11,33,34], RNA-sequencing (RNA-seq)[35], and assay for transposase-accessible chromatin using sequencing (ATAC-seq)[36]. In this study, we exploit these modern approaches to assess the suitability of SN4741 as an *in vitro* proxy for DA neurons and determine the extent to which this cell line is appropriate for prioritizing and investigating the mechanisms by which PD-associated variation confers disease risk.

Through a combination of karyotyping, single-cell (sc)RNA-seq, and RT-qPCR, we evaluate the genomic integrity of this immortalized cell line, determine how the transcriptional profile and expression of DA neuron marker genes in this line changes between undifferentiated (37°C) and differentiated (39°C) states, and evaluate whether these transcriptional changes are consistent throughout the differentiation process. The data we collect suggests that while these cells show evidence that they are exiting a proliferative state and entering a more differentiated state, they are an unsuitable model of SN DA neurons, as they possess aneuploidy and structural abnormalities, as well as consistently low expression of DA neuron markers upon differentiation. We employ bulk RNA-seq to quantify transcriptional differences between differentiated and undifferentiated SN4741 cells and determine that, while transcriptional profiles change to reflect differentiation, they do not show strong evidence that these cells are entering a DA state. We then compare chromatin accessibility profiles of undifferentiated and differentiated SN4741 cells with those of *ex vivo* mouse E15.5 midbrain (MB) and forebrain (FB) neurons and determine that the chromatin accessibility profiles of SN4741 cells do not reflect the cellular population from which they were derived. Collectively, cytogenetic, chromatin, and transcriptional data suggest that the SN4741 cell line is not as strong a cellular surrogate for DA neurons as previously thought. Ultimately, this work underscores the importance of leveraging technological advances in genomic and cellular analyses to evaluate, and re-evaluate, the suitability of established model systems in disease biology.

## RESULTS

### SN4741 is an Unstable Polyploid Cell Line

G-band karyotyping was performed on 20 SN4741 metaphase spreads and a representative karyogram (**Figure 1A**) was generated. The karyotype was interpreted as an abnormal, polyploid, karyotype with complex numerical abnormalities and unbalanced, structural abnormalities. While most, but not all, abnormalities were consistently present in these cells; none of the 20 cells assessed had the same chromosome complement, and no normal cells were observed. All cells possessed at least one copy of each mouse autosome (1 through 19) and female sex chromosomes; however, most chromosomes were triploid in each cell (**Figure 1B**). These karyotypic abnormalities already call into question the viability of these cells as a surrogate for human neurodegenerative disease. Since these cells are genetically unstable, there may be large experimental batch effects as the cell populations shift across divisions. Furthermore, gene dosage effects that severely deviate from normal copy number in DA neurons may lead to confounding and unreliable results.

**Figure 1.**
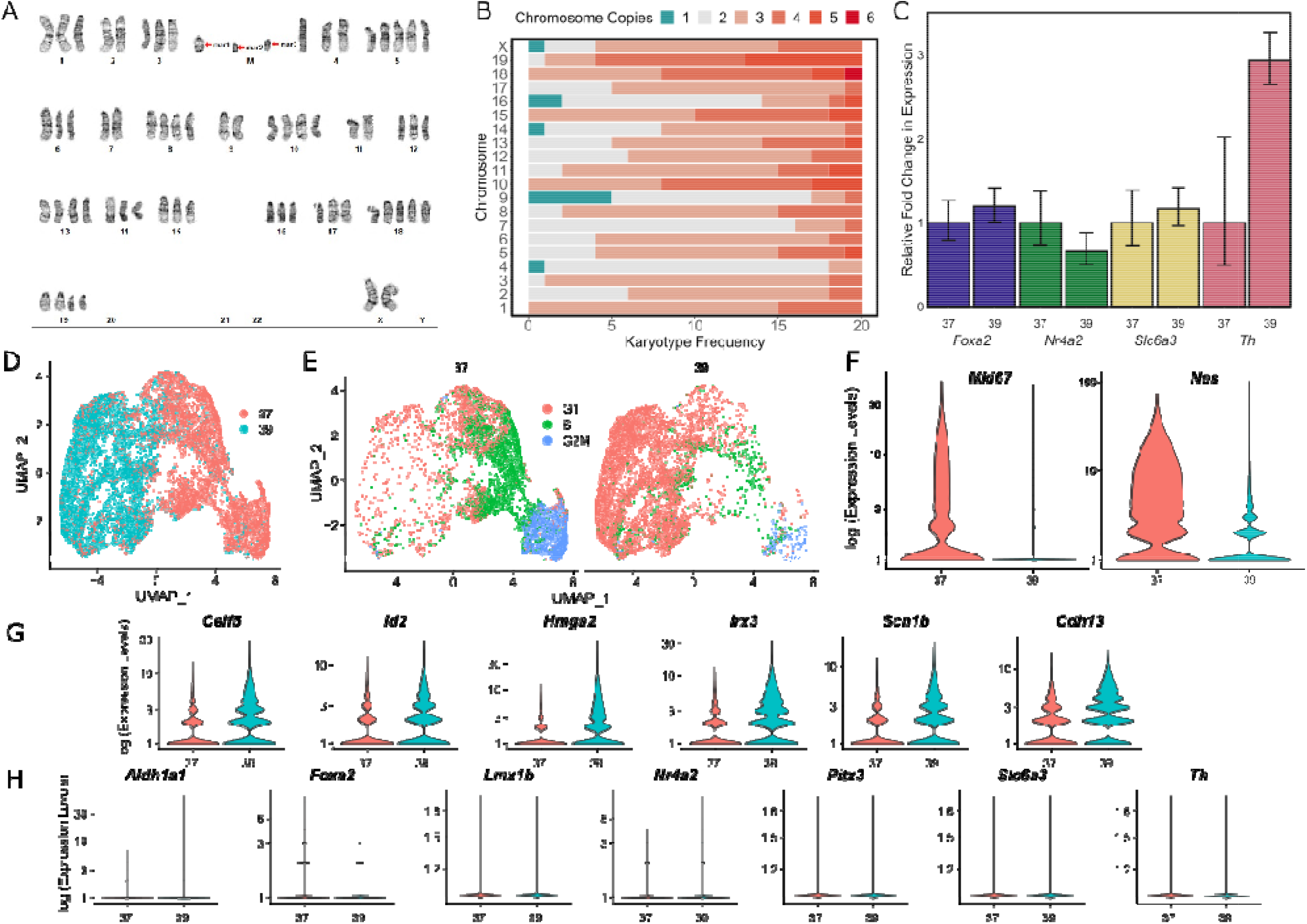
Characterizing the genomic stability and differentiation consistency of the temperature sensitive SN4741 cell line. **A**) A representative karyogram of SN4741 cells, indicating structural instability (M; marker chromosomes) and unstable triploidy. **B**) A stacked bar plot summarizing the aneuploidy frequency of each chromosome over 20 SN4741 karyotypes. **C**) Assaying expression of dopaminergic neuron markers by RT-qPCR indicates that *Foxa2, Nr4a2, and Slc6a3* remain at similar expression levels, but *Th* expression increases, when SN4741 cells are shifted from the permissive temperature (37°C) to the non-permissive temperature (39°C). **D**) UMAP plot of scRNA-seq at the permissive and non-permissive temperatures indicates that cells at each temperature are transcriptionally distinct. **E**) Analysis of scRNA-seq data demonstrates that shifting the cells to the non-permissive temperature is accompanied by a shift in cell cycle stage from G2M and S phases to primarily G1 phase. **F**) Violin plots generated with scRNA-seq data show that *Mki67*, a marker of cellular proliferation, and Nes, a neural stem cell marker, are both expressed at the permissive temperature (37°C), with little to no expression at the non-permissive temperature (39°C). **G**) Violin plots generated with scRNA-seq data show that transcripts associated with immature neurons are upregulated when SN4741 cells are shifted to the non-permissive temperature. **H**) Violin plots generated with scRNA-seq data show that expression of DA neural markers, *Aldh1a1, Foxa2, Lmx1b, Nr4a2, Pitx3, Slc6a3, and Th*, remain at similar levels when SN4741 cells are shifted to the non-permissive temperature.

### Undifferentiated and Differentiated SN4741 Cells Express Similar Levels of Dopaminergic Neuron Marker Genes by RT-qPCR

Preliminary expression analysis by RT-qPCR confirmed expression of a variety of DA neuron markers: forkhead box A2 *(Foxa2)*, nuclear receptor subfamily 4 group A, member 2 (*Nr4a2)*, solute carrier family 6 member 3 *(Slc6a3)*, and tyrosine hydroxylase *(Th)*. Compared to the expression of these markers in the undifferentiated SN4741 cell culture (37°C), relative expression of all markers remained at similar levels when the cells were shifted to the higher temperature condition (39°C), except for an increase in *Th* expression (**Figure 1C**). While elevated *Th* has been used as a marker of differentiation into DA neurons in previous work with SN4741 cells[20,37,38], an increase in *Th* expression is not exclusively associated with DA neurons. *Th* is a marker for all catecholaminergic neurons (dopaminergic and adrenergic)[39], and evidence suggests that *Th* expression is transient in other neurons throughout embryonic development[40–42]. These results indicate that at the non-permissive temperature, SN4741 cells may not be fully differentiating into DA neuron progenitors.

### scRNA-seq Reveals that SN4741 Cells Differentiate at the Non-Permissive Temperature, but Lack Expression of DA Neuron Marker Genes

To assess the consistency of the differentiation protocol, transcriptomes were generated from ≥17,000 cells across four replicates cultured at the permissive temperature (37°C) and four replicates cultured at the non-permissive temperature (39°C). Analysis of the single-cell transcriptomes reveal that the cells cluster by growth temperature (**Figure 1D**). This separation of cells by temperature is accompanied by changes to the cell cycle, with cells at the permissive (37°C) temperature mostly in either G2M or S phase, while cells at the non-permissive temperature (39°C) are mostly in G1 phase (**Figure 1E**), indicating that they may be differentiated. In expression analysis, markers of proliferation that are expressed in G2M phase, like Marker of Proliferation Ki-67 (*Mki67*), are predominantly expressed in cells at the permissive temperature (**Figure 1F**), corroborating the cell cycle analysis. When shifted to the non-permissive temperature, SN4741 cells appear to robustly differentiate, exemplified by a decrease in the expression of Nestin (*Nes*), a neural stem cell marker (**Figure 1F**). Additional transcriptional changes at this non-permissive temperature include an increase in the expression of a neural marker CUGBP Elav-Like Family Member 5 (*Celf5*)[43], as well as genes that have been found to regulate neural stem cell self-renewal (Inhibitor of DNA Binding 2, *Id2;* High Mobility Group AT-Hook 2, *Hmga2*)[44,45], neurogenesis (Iroquois Homeobox 3, *Irx3*)[46], and arborization of neurons (Sodium Voltage-Gated Channel Beta Subunit 1, *Scn1b*)[47], indicating that these cells may be differentiating into neural precursor cells (**Figure 1G**). Furthermore, Cadherin 13 (*Cdh13)*, a modulator of GABAergic neurons, is significantly upregulated at this non-permissive temperature (**Figure 1G**), while the expression of a variety of DA neuron markers fail to be detected in either the permissive or non-permissive temperatures. Markers, including Aldehyde Dehydrogenase 1 Family Member A1 (*Aldh1a1), Foxa2*, LIM Homeobox Transcription Factor 1 Beta *(Lmx1b), Nr4a2*, Paired-like homeodomain 3 (*Pitx3), Slc6a3*, and *Th* have few to no reads assigned to them **(Figure 1H)**. Collectively, these results suggest that while SN4741 cells are differentiating towards a neuronal fate when shifted to the nonpermissive temperature, they may not be entering a clear DA trajectory under these conditions.

### ATAC-seq Identifies Differential Open Chromatin Profiles in SN4741 Cells at the Permissive and Non-Permissive Temperatures

To consider how chromatin accessibility changes between the two temperatures, we performed ATAC-seq on SN4741 cells in both the undifferentiated and differentiated states. Libraries were confirmed to be technically and biologically relevant (**Supplemental Figure 1**), and well correlated between replicates (**Supplemental Figure 2; Figure 2A-B**).

**Figure 2:**
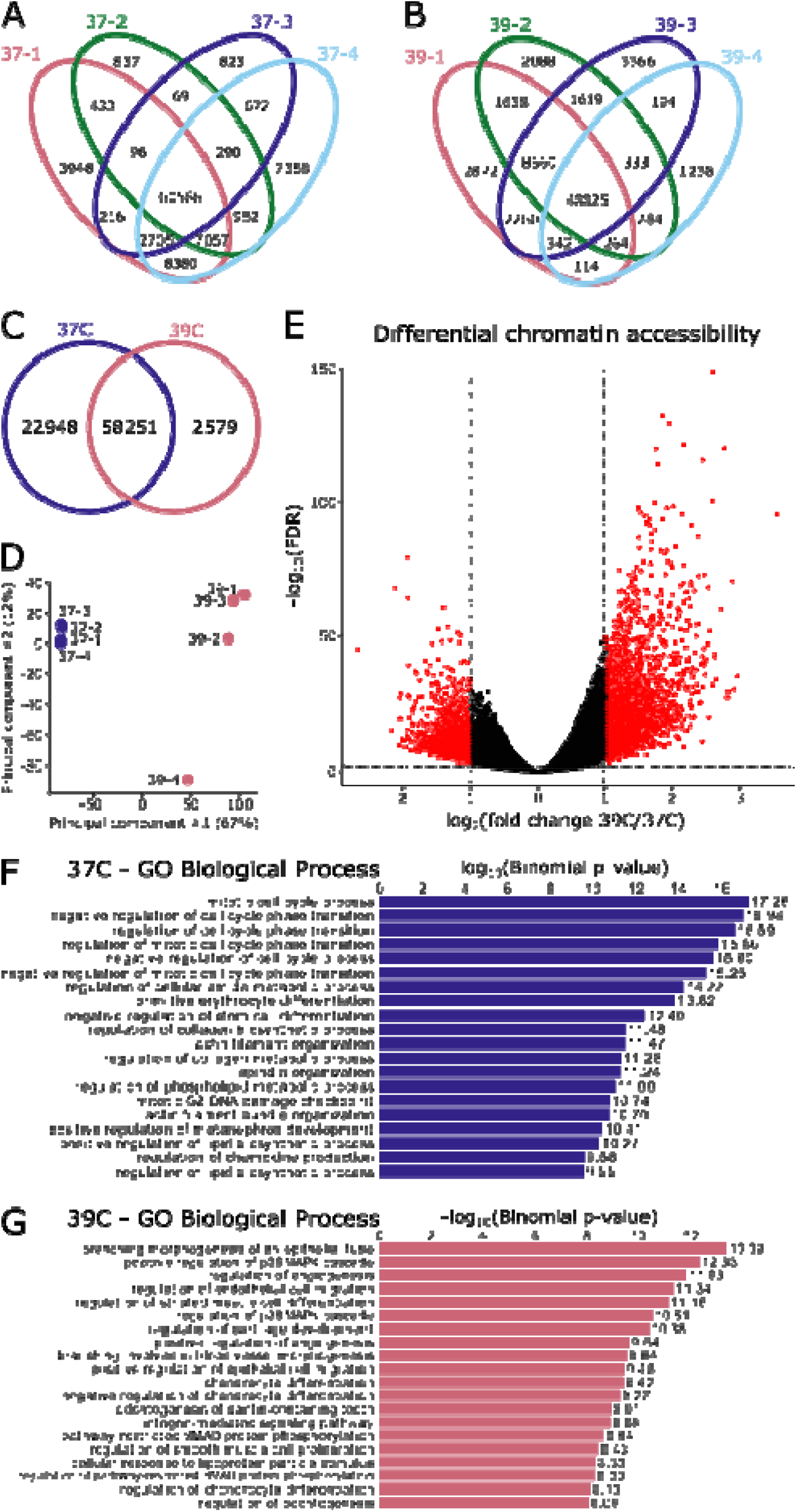
Changes in chromatin accessibility suggest a reduction in potency at the non-permissive temperature. **A, B)** Replicates are highly similar within temperature conditions, with the majority of peaks present in all four replicates. **C**) The two temperatures share 58,251 regions of open chromatin but do not overlap completely. **D**) Principal component analysis resolves the two temperatures on the first principal component. **E**)Differential accessibility analysisidentifies 5,055 differentially accessible regions, with 2,654 preferentially open in the permissive temperature (37°C) and 2,401 preferentially accessible at the non-permissive temperature (39°C). **F**) Gene ontology (GO) of genes adjacent regions that are preferentially open at the permissive temperature are associated with regulation of the cell cycle and negative regulation of differentiation, as is appropriate for this temperature. **G**) Gene ontology of genes adjacent regions that are preferentially open at the non-permissive temperature are associated with a variety of differentiation fates (blood vessels, muscle cells, cartilage/chondrocytes). Additionally, two of the top gene ontology terms relate to the p38 MAPK cascade, which has been found to be activated as a cellular response to heat stress.

A total of 83,778 consensus open chromatin regions were identified, with 70% of peaks shared between the two temperatures (**Figure 2C**). Principal component analysis of these consensus regions suggests a clear separation in the chromatin state between the two temperatures (**Figure 2D**). To explore these differences, we performed differential accessibility analysis with DiffBind[48], to find a total of 5,055 differentially accessible regions: 2,654 enriched in the permissive temperature and 2,401 enriched at the non-permissive temperature (log_2_FC > 1, FDR < 0.05; **Figure 2E**).

Gene ontology of genes adjacent to differentially accessible regions largely recapitulate the scRNA-seq analysis; functions associated with regions preferentially open at the permissive temperature suggest the maintenance of the undifferentiated, cell-cycling state (**Figure 2F**). The gene ontology of genes adjacent to those regions preferentially accessible at the non-permissive temperature is less coherent and suggest cell differentiation towards several fates (blood vessels, cartilage, tooth), none of which are neuronal and, perhaps unsurprisingly, demonstrate evidence of response to temperature stress[49] (**Figure 2G**).

Overall, there is a shift in the chromatin accessibility between the two temperatures that indicate the cells transition from an undifferentiated to differentiated state as the cells move from the permissive to non-permissive temperature. The differences in chromatin accessibility further confirm that SN4741 cells are not differentiating towards a neuronal lineage.

### Comparison of chromatin accessibility in SN4741 cells fails to recapitulate the chromatin landscape of *ex vivo* mouse DA neurons

To evaluate the potential relationship between SN4741 cells and DA neurons they are presumably modelling, we compared the chromatin accessibility between the SN4741 cells at both temperatures to previously generated *ex vivo* mouse embryonic DA neuron chromatin accessibility profiles (NCBI GEO: GSE122450;[50]).

Considering the consensus peak set of 165,334 regions generated from all *in vivo* and *ex vivo* samples and their normalized read counts, we observe a clear separation between the SN4741 cell culture model and the *ex vivo* DA neurons by correlation and principal component analysis (**Figure 3A, B**). Examining the raw overlap of peaks between the SN4741 cells and *ex vivo* neurons, just 12.5% (20,667) are present in all four cell types/conditions (**Figure 3C**). The chromatin profiles are largely exclusive between the SN4741 cell culture model and the *ex vivo* DA neurons: 41.3% (68,304) of regions are accessible solely in the *ex vivo* neuron populations and 40% (65,857) are exclusively accessible in the SN4741 cell culture models. There is little overlap between the *ex vivo* and cultured samples. In comparison of the *ex vivo* midbrain DA neurons to the non-permissive, differentiated temperature, only 183 peaks are restricted to these populations.

**Figure 3:**
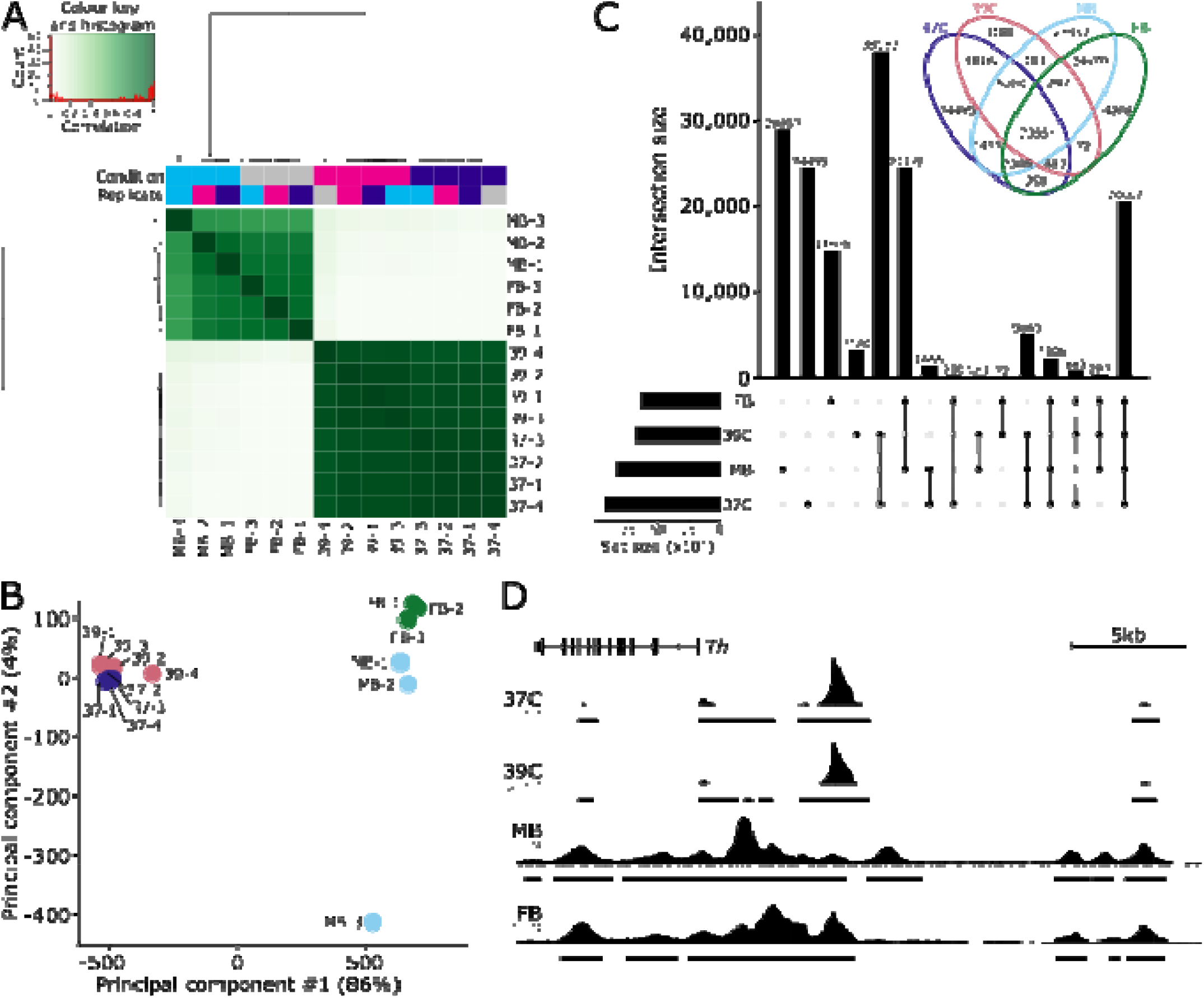
Chromatin accessibility of SN4741 cells do not resemble *ex vivo* dopaminergic neurons. **A)** SN4741 samples are highly correlated with each other but very poorly correlate with the open chromatin landscape of either midbrain (MB) or forebrain (FB) embryonic mouse dopaminergic neurons. **B**) Principal component analysis shows a clear separation between the *ex vivo* and *in vitro* samples along PC1, representing 86% of the variance. **C**) An upset plot and associated Venn diagram quantify the overlap of peaks between the four conditions and show the poor relationship between the SN4741 cells and the *ex vivo* mouse dopaminergic neurons. Most peaks are specific to a single cell type/temperature or are restricted to either the *ex vivo* or *in vitro* samples. Few peaks are specifically shared between the non-permissive temperature and the *ex vivo* samples; for example, there are just 183 peaks that are shared exclusively by the MB dopaminergic neurons and the SN4741 cells at the non-permissive temperature. **D**) A genome track showing the normalized read pile up and called consensus peaks in each of the cell types/temperatures at the key dopaminergic neuron specification gene, *Th*. The chromatin accessibility is largely similar within *ex vivo* or *in vitro* cells but bear little resemblance to each other.

The chromatin profiles between *ex vivo* embryonic DA neurons and their prospective *in vitro* cell culture surrogate are virtually independent. They exhibit scant overlap in their global chromatin profiles and bear little resemblance to each other at regulatory regions of key DA neuron genes (**Figure 3D**). Neither the analysis of the SN4741 chromatin accessibility profiles in isolation or in comparison with *ex vivo* neurons would suggest these cells to be appropriate models of embryonic DA neurons.

### Transcriptional Changes in SN4741 Cells Indicate Differentiation from Pluripotent Stem Cells into Brain Cells that do not Fully Resemble MB DA Neurons

Bulk RNA-seq data were also generated for SN4741 cells, at both the permissive and non-permissive temperatures, to determine whether transcriptome changes reflect differentiation into DA neurons, or other neural cell types. To evaluate the RNA-seq libraries, quality-control measures were performed *in silico* **(Supplemental Figure 3)**. PCA **(Supplemental Figure 3B)**, and sample-sample distances **(Supplemental Figure 3C**) reaffirmed that samples cultured at the same temperature are more like one-another than samples cultured at the alternate temperature.

We found that 735 genes were upregulated at the non-permissive temperature (adjusted p-value < 0.01 and log_2_ FC > 1.5), and 954 genes were downregulated (adjusted p-value < 0.01 and log_2_ FC< -1.5) at the non-permissive temperature. The list of genes significantly downregulated at the non-permissive temperature was submitted to Enrichr (https://maayanlab.cloud/Enrichr/) [51–53] for gene ontology (GO) and analysis of cell type markers. Consistent with the observation that cells at the non-permissive temperature are differentiated and in G1 phase of the cell cycle, downregulated genes resulted in GO terms strongly enriched for mitotic and DNA replication processes (**Figure 4A)**. Additionally, significantly downregulated genes at the non-permissive temperature overlap with subsets of PanglaoDB[54] cell type marker genes, suggesting that these cells are shifting away from a state that resembles neural stem cells (**Figure 4B**).

**Figure 4.**
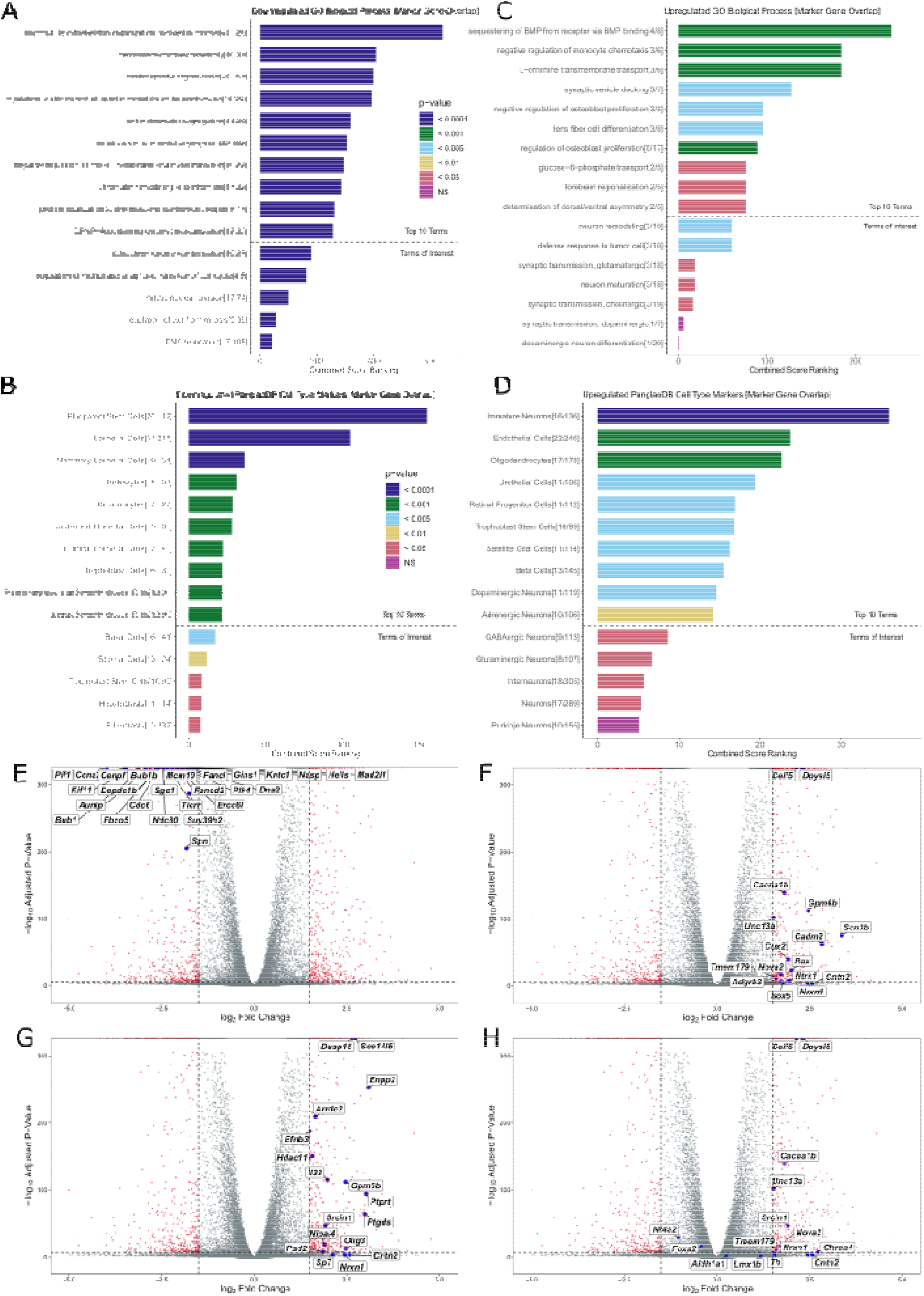
Gene Ontology and Differential Expression Analysis of Bulk RNA-seq Data: **A)** Top 10 GO terms for downregulated DE genes in SN4741 cells at the non-permissive temperature, followed by GO terms of interest (below dotted line). Terms were evaluated using Combined Score Ranking = (p-value computed using the Fisher exact test)*(z-score computed by assessing the deviation from the expected rank), based on enrichment of DE genes that overlap with Enrichr input genes for each term (the end of each bar). **B**) Top 10 predicted cell types based on downregulated DE genes in SN4741 cells at the non-permissive temperature, followed by predicted cell types of interest (below dotted line). Terms were evaluated using Combined Score Ranking = (p-value computed using the Fisher exact test)*(z-score computed by assessing the deviation from the expected rank), based on enrichment of DE genes that overlap with PanglaoDB input genes for each term (the end of each bar). **C**) Top 10 GO terms for downregulated DE genes in SN4741 cells at the non-permissive temperature. **D**) Top 10 predicted cell types based on upregulated DE genes in SN4741 cells at the non-permissive temperature. **E**) Volcano plot of –log10 adjusted p-value versus log2 fold change with DESeq2 after lfc shrinkage, contrasting the fold change in expression of SN4741 cells at 39°C, using SN4741 cells at 37°C as reference. Red points = genes that are statistically differentially expressed (adjusted p-value < 0.01, |log2FC| > 1.5). Blue points = Overlapping immature neuron marker genes. **F**) Blue points = Overlapping immature neuron marker genes **G**) Blue points = Overlapping oligodendrocyte marker genes. **H**) Blue points = Overlapping DA neuron marker genes.

Similarly, the list of significantly upregulated genes was submitted to Enrichr for GO and analysis of cell type markers. As expected, upregulated genes resulted in GO terms for biological processes that indicate a more terminally differentiated cell type **(Figure 4C**): “synaptic vesicle docking,” “negative regulation of osteoblast proliferation,” “lens fiber cell differentiation,” “regulation of osteoblast proliferation,” and “forebrain regionalization”. While not included in the top 10 terms by combined score ranking, “neuron remodeling,” “synaptic transmission, glutamatergic,” “neuron maturation,” and “synaptic transmission, cholinergic” were also identified as significantly associated terms. Notably, “synaptic transmission, dopaminergic” and “dopaminergic neuron differentiation” were also listed as insignificant terms (**Figure 4C**), as *Th* was the lone overlapping marker gene for these terms.

In line with GO terms enriched for biological processes involving differentiation, possibly in neuronal cells, overlapping PanglaoDB[54] cell type marker genes suggest that SN4741 cells at the non-permissive temperature most significantly resemble immature neurons (**Figure 4D**). “Oligodendrocytes,” “retinal progenitor cells,” “satellite glial cells,” “dopaminergic neurons,” “adrenergic neurons,” “GABAergic neurons,” and “glutamatergic neurons” were also listed as cell types with significant marker gene overlap.

The distribution of various cell type marker genes on a volcano plot, indicating the log_2_FC in expression and -log_10_ adjusted p-values of DE genes, reveals that the specific genes overlapping “pluripotent stem cell” markers (26/112), cluster as the most highly significantly downregulated genes **(Figure 4E)**. In contrast, only two of the upregulated marker genes overlapping “immature neurons” (16/136, **Figure 4F**) and “oligodendrocytes” (17/178, **Figure 4G**) cluster in a similarly strong way. Plotting the 11/119 overlapping upregulated genes for “dopaminergic neurons” (**Figure 4H**) reveals that 7/11 overlapping genes (*Celf5, Dpys15, Cacna1b, Tmem179, Nova2, Nrx1*, and *Cntn2*) are also marker genes for immature neurons. Plotting the DA neuron markers also assayed by RT-qPCR validates that the relative expression of these markers is consistent between these highly sensitive assays. At 39°C, *Th* expression increases (log_2_FC = 1.552651); *Nr4a2* expression decreases (log_2_FC = -1.042794), and expression of *Slc6a3* and *Foxa2* is not significantly different between conditions: (*Slc6a3* was filtered out due to low read counts across both temperature conditions and *Foxa2* log_2_FC = - 0.4142541).

To confirm the GO-indicted cell types, normalized read counts for select marker genes were plotted for each temperature replicate: *Celf5*[43], *Nrxn1*[55], *Ntrk1*[56], and *Unc13a*[57], for “immature neurons” were upregulated at 39°C relative to cells at 37°C (**Figure 5A**); *Olig3*[58], *Il33*[59], *Hdac11*[60], and *Ptgds*[61] for “oligodendrocytes” were upregulated at 39°C relative to cells at 37°C (**Figure 5B**); and *Ccna2*[62], *Cdc6*[63], *Cenpf*[64], and *Gins1*[65] for “pluripotent stem cells” were downregulated at 39°C relative to cells at 37°C (**Figure 5C**).

**Figure 5.**
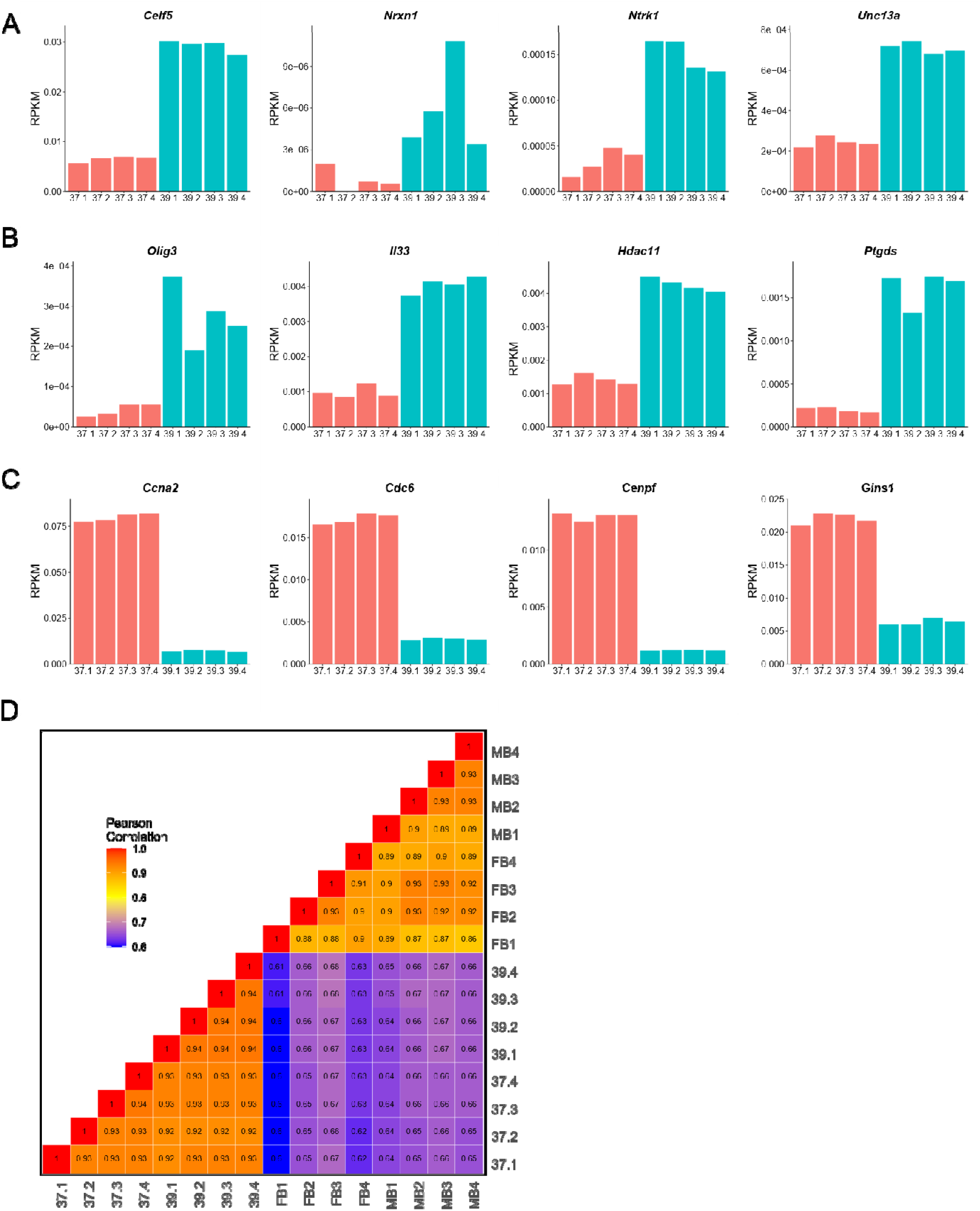
Validation of gene ontology and comparison to bulk-RNA-seq from ex vivo DA neurons. **A)** A bar chart showing normalized bulk RNA-seq read counts from genes upregulated at 39°C that overlap the “immature neurons” predicted cell type. **B**) A bar chart showing normalized bulk RNA-seq read counts from genes upregulated at 39°C that overlap the “oligodendrocyte” predicted cell type. **C**) A bar chart showing normalized bulk RNA-seq read counts from genes downregulated at 39°C that overlap the “pluripotent stem cell” predicted cell type. **D**) A Pearson correlation heatmap comparing the transcriptomes of SN4741 cells at 37°C and 39°C to midbrain (MB) or forebrain (FB) embryonic mouse dopaminergic neurons.

The differentially expressed gene sets were then analyzed using STRING (https://string-db.org/)[66]. The set of upregulated genes was enriched for protein-protein interactions (number of edges = 705; expected number of edges = 438; PPI enrichment p-value = < 1.0e-16) and GO terms such as “Neuron differentiation” (GO:0030182; FDR = 0.0016), “Neuron development” (GO:0048666; FDR = 0.0070), and “Neurogenesis” (GO:0022008; FDR = 0.0085) supporting that this upregulated gene set is a meaningful group likely belonging to a network involved in neuronal maturation. The set of downregulated genes was also enriched for protein-protein interactions (number of edges = 9251; expected number of edges = 1910; PPI enrichment p-value = < 1.0e-16) and GO terms such as “Cell cycle” (GO:0007049; FDR = 5.90e-36), “Mitotic cell cycle” (GO:0000278; FDR = 6.97e-34), and “Cell division” (GO:0051301; 6.35e-25) further confirming that these cells are no longer undergoing cell division.

Finally, previously generated reads per kilobase of exon per million reads mapped (RPKM) from *ex vivo* E15.5 mouse embryonic DA neuron bulk RNA-seq[50] were used to compare how closely the SN4741 transcriptome resembles the neuronal populations they are expected to model. Similar to our results comparing chromatin accessibility between these two datasets, correlation of RPKM shows a clear separation between the SN4741 cell culture model and the *ex vivo* DA neurons (**Figure 5D**). Collectively, these results confirm that at the non-permissive temperature, SN4741 cells are no longer rapidly dividing, neural stem cells. However, while the transcriptional profile of these cells indicates that they are differentiating towards cell types present in the brain, these cells do not fully possess characteristics of the MB DA neurons they are meant to model.

## DISCUSSION

It is critically important that studies of human disease generate biologically accurate data, whether aimed at elucidating molecular mechanisms, onset and progression, or management and therapeutics. In the context of discovery biology or the illumination of human health and disease mechanisms, misattribution of cellular identity, or other deviations from biological accuracy, may result the misinterpretation of biological findings or misdirected research efforts. When studying human disease, cellular surrogates are often used to overcome the ethical and technical limitations of employing animal models. Therefore, it is imperative that disease-relevant insights are predicated on robust data generated from model systems representing human biology as accurately as possible.

Here, we demonstrate the importance of assessing *in vitro* models of disease to determine the extent to which they can yield biologically accurate data that can be used to inform aspects of human disease. The SN4741 cell line has been used to study neurotoxicity and therapeutic interventions [21–27], PD-associated genetic mutation[28,29] and cell signaling and transcriptional regulation[30–32,37], since it was initially characterized as an immortal, mouse MB-derived cell line that differentiates into DA neurons at a non-permissive temperature[20]. However, contemporary genomic analyses have not been leveraged to characterize and evaluate the SN4741 cell line as a suitable proxy for DA neurons in PD, until now.

We employed karyotyping, RT-qPCR, and scRNA-seq to assess the genomic stability of these cells and determine how consistently they differentiate into DA neurons at the non-permissive temperature. We generated bulk RNA-seq and ATAC-seq data from this cell line at both the permissive and non-permissive temperatures, to extensively characterize this cell line and document how transcriptional landscapes and chromatin accessibility profiles shift in response to temperature-induced differentiation and compare to known profiles of *ex vivo* DA neurons. Our results suggest that SN4741 is an unstable, polyploid cell line that is unlikely to be a viable differentiation model of DA neurons; and thus, is likely not a robust proxy by which to study MB DA neurons in the context of human phenotypes, including PD, schizophrenia, addiction, memory, or movement disorders.

The results of karyotyping alone indicate that any data generated using SN4741 cells may be biologically inaccurate due to extreme variability in chromosome complement and therefore, copy number variation, between individual cells. Consequently, the results of previous studies evaluating neurotoxicity[21,23,24,26,27,30,32,38,67], cellular signaling pathways[22,25,28,68,69], and transcriptional profiling[37] in these cells may have been unduly influenced by the extreme imbalance in gene dosage that we found to vary from cell to cell. For example, alpha-synuclein (*SNCA)* has been consistently implicated in PD risk[70–73], particularly due to variants that promote α-synuclein misfolding[74] and overexpression[50] or events that result in gene amplification [75,76]. *Snca* is present on mouse chromosome 6 and the karyotypes generated for SN4741 cells show that chromosome 6 is triploid in most assayed cells (**Figure 1B**). Therefore, using the SN4741 cell line to model neurodegeneration in PD may result in inaccurate data due to an exaggerated vulnerability towards degeneration imposed by elevated *Snca* copy number, by gene dosage effects of other interacting gene products in relevant pathways, or by the structural instability of this line.

Even if this cell line could be adopted to study *Snca* overexpression/amplification, ATAC-seq profiling of open chromatin regions in this cell line at the permissive and non-permissive temperatures indicates that these cells do not possess chromatin accessibility profiles similar to those of *ex vivo*, mouse E15.5 MB neurons. In PD, disease is characterized by the degeneration of MB DA neurons, while DA neurons of the FB are spared. Therefore, the chromatin profiles of MB DA neurons, as well as the differentially accessible regions of the genome between MB and FB neurons, may influence the preferential vulnerability of MB neurons in PD[50]. In the context of exploiting these chromatin profiles to study PD-associated variability and neurotoxicity, SN4741 cells are likely a poor model, as the open chromatin regions of these cells are not a reliable proxy for mouse E15.5 MB or FB DA neurons.

The chromatin accessibility profiles of SN4741 cells not only fail to cluster with *ex vivo* populations of mouse MB neurons, but the transcriptional landscapes of these cells suggest that these cells have shifted towards a more differentiated state that may be less DA than previously thought. Examination of cell cycle markers by scRNA-seq demonstrates that SN4741 cells at the non-permissive temperature are more differentiated than cells at the permissive temperature, as expected[20]. GO terms for genes that are significantly downregulated at the non-permissive temperature reinforces that these cells are no longer rapidly dividing, pluripotent stem cells. However, RT-qPCR, scRNA-seq, and bulk RNA-seq in these cells fail to detect significant upregulation of most key DA neuron markers in the differentiated cells, except for *Th. Th* is not exclusively expressed by DA neurons at embryonic timepoints[40–42]. In fact, significantly upregulated genes in SN4741 cells at the non-permissive temperature that overlap with GO terms and cell cycle marker genes suggests that *Th* is the only significantly upregulated gene overlapping with biological processes involving DA neurons. Rather, additional overlapping cell type marker genes suggest that these cells more closely resemble immature neurons.

In parallel, we generated promoter capture (pc)Hi-C data at the non-permissive temperature with the intention of exploring how non-coding disease-relevant variants interact with promoters and potentially regulate gene expression in MD DA neurons. As our group is focused on PD-associated variation, which is unlikely to act broadly in immature neurons, our group has not analyzed the resulting data, beyond basic quality control (**Supplemental Figure 4**). While the SN4741 cells at the non-permissive temperature fail to recapitulate the transcriptomic or chromatin state of DA neurons, it is of potential interest for follow-up studies that they do resemble some immature neuron types. Although not analyzed by our group, we generated output files for interaction detection, and this data has been made publicly available for others to explore (accessible through: https://github.com/rachelboyd/SN4741_pcHiC), as it may be useful to study genomic interactions at promoters driving an immature neuronal state. However, the cell type best represented by SN4741 cells at the non-permissive temperature still requires deeper characterization.

Any data generated using SN4741 cells in the context of DA neuron modeling and/or PD must be interpreted with caution and in light of the appropriate caveats. Due to the instability and polyploidy of this cell line, we recommend that the use of SN4741 cells for PD-related research be re-evaluated. Future studies designed to fine-tune the classification of these cells may support the use of SN4741 cells as a model of other neuronal or non-neuronal cells. Additionally, the differentiation trajectory of these cells may be amenable to intervention(s) that could drive their molecular state towards one that resembles DA neurons more closely.

## CONCLUSIONS

This study establishes a valuable precedent with broad implications across biological and disease-related research. Prior to using SN4741 cells to study non-coding regulatory variation in PD, we characterized this cell line to determine its suitability as a model of DA neurons in PD, and found that these cells are unstable, polyploid cells that do not demonstrate strong molecular characteristics of MB DA neurons. These cells express low levels of DA neuron markers, and chromatin landscapes in differentiated SN4741 cells scarcely overlap open chromatin regions in *ex vivo* mouse E15.5 midbrain neurons. We demonstrate the importance of genomic characterization of *in vitro* model systems prior to generating data and valuable resources that may be used to inform aspects of human disease. In future studies that utilize *in vitro* models of any human disease, due diligence to confirm their suitability as surrogates could save time, resources, and possibly lives, by avoiding misdirection and advancing successful therapeutic development.

## METHODS

### Cell Culture

SN4741 cells were obtained from the Ernest Arenas group at the Karolinska Institutet. SN4741 cells were confirmed to be mycoplasma free using a MycoAlert® Mycoplasma Detection Assay (Lonza) and were maintained in high glucose Dulbecco’s Modified Eagle Medium (DMEM; Gibco 1196502), supplemented with 1% penicillin–streptomycin and 10% fetal bovine serum (FBS) in a humidified 5% CO_2_ incubator at 37°C. Cells at 80% confluence were passaged by trypsinization approximately every 2-3 days. To induce differentiation, 24 hours after the cells were passaged, media was replaced by DMEM supplemented with 1% penicillin–streptomycin and 0.5% FBS at 39°C. Cells were allowed to grow and differentiate in these conditions for 48 hours before harvesting for experimentation.

### G-Band Karyotyping

At passage 21, undifferentiated SN4741 cells were sent to the WiCell Research Institute (Madison, Wisconsin), at 40-60% confluency, for chromosomal G-band analyses. Karyotyping was conducted on 20 metaphase spreads, at a band resolution of >230, according to the International System for Human Cytogenetic Nomenclature.

### cDNA Synthesis and RT-qPCR for DA neuron markers

RNA was extracted from both differentiated and undifferentiated cells by following the RNeasy Mini Kit (QIAGEN) protocol, as written. 1 μg of each RNA sample underwent first-strand cDNA synthesis using the SuperScript III First-Strand Synthesis System for RT-PCR (Invitrogen) according to the oligo(dT) method. qPCR was performed with Power SYBR Green Master Mix (Applied Biosystems), using primers for β-actin (*Actb*), *Foxa2, Nr4a2, Slc6a3*, and *Th* (**Table S1**). Reactions were run in triplicate under default SYBR Green Standard cycle specifications on the Viia7 Real-Time PCR System (Applied Biosystems). Normalized relative quantification and error propagation followed the data analysis and associated calculations proposed by Taylor *et al*. (2019)[77], with results normalized to *Actb*.

### Single cell RNA-seq Library Preparation, Sequencing, and Alignment

Both differentiated (39°C) and undifferentiated (37°C) cells were trypsinized, and scRNA-seq libraries were generated following the Chromium 10X pipeline[78]. Four replicates at each temperature across >17,000 cells were assayed. Cell capture, cDNA generation, and library preparation were performed with the standard protocol for the Chromium Single Cell 3’ V3 reagent kit. Libraries were quantified with the Qubit dsDNA High Sensitivity Assay (Invitrogen) in combination with the High Sensitivity DNA Assay (Agilent) on the Agilent 2100 Bioanalyzer. Single-cell RNA-sequencing libraries were pooled and sequenced on an Illumina NovaSeq 6000 (SP flow cells), using 2×50 bp reads per library, to a combined depth of 1.6 billion reads. The quality of sequencing was evaluated via FastQC. Paired-end reads were aligned to the mouse reference genome (mm10) using the CellRanger v3.0.1 pipeline. Unique molecular identifier (UMI) counts were quantified per gene per cell (“cellranger count”) and aggregated (“cellranger aggr”) across samples with no normalization.

### Single Cell RNA-seq Analysis

Using Seurat[79](v4.2.0), cells were filtered to remove stressed/dying cells (% of reads mapping to the mitochondria > 15%) and empty droplets or doublets (number of unique genes detected <200 or >6,000). Cells were scored for their stage in the cell cycle using “CellCycleScoring()” on cell cycle genes provided by Seurat (“cc.genes”). Cells were then normalized using “SCTransform” (vst.flavor = “v2”) and corrected for percent mitochondrial reads and sequence depth. Principal component (PC) analysis was performed and a PC cut-off was identified using “ElbowPlot().” Using this PC cutoff and a minimum distance of 0.001, UMAP clustering was used for dimensionality reduction. Expression was plotted on a log scale with “VlnPlot()” for a variety of proliferation and DA neuron markers.

### ATAC-seq Library Preparation and Quantification

ATAC-seq libraries were generated for four replicates of undifferentiated (37°C) and differentiated (39°C) SN4741 cells, according to the Omni-ATAC protocol[80], with minor modifications. Aliquots of 50,000 cells were centrifuged at 2000 x g for 20 mins at 4°C, and the resulting pellets were resuspended in 50μL of resuspension buffer. Cells were left to lyse for 3 minutes on ice before being centrifuged again at 2000 x g for 20 minutes at 4°C. The resulting nuclei pellets were then tagmented, as written, using 50μL of transposition mixture and then incubated at 37°C for 30 mins in a 1000 RPM thermomixer. After transposition, DNA was purified with the Zymo DNA Clean and Concentrator -5 Kit and eluted in 21μL of elution buffer. Pre-amplification of the transposed fragments was performed according to the conditions outlined in the Omni-ATAC protocol[80]; however, 12 pre-amplification cycles were run in lieu of qPCR amplification to determine additional cycles. The amplified libraries were prepared according to the Nextera DNA Library Prep Protocol Guide, except that libraries were purified with 40.5μL AMPure XP beads (Beckman Coulter), and 27.5μL of resuspension buffer was added to each sample. All libraries were quantified with the Qubit dsDNA High Sensitivity Assay (Invitrogen) in combination with the High Sensitivity DNA Assay (Agilent) on the Agilent 2100 Bioanalyzer.

### ATAC-seq Sequencing, Alignment, and Peak Calling

Libraries were sequenced on Illumina NovaSeq 6000 (SP flow cells), using 2×50 bp reads per library, to a total combined depth of 1.6 billion reads. The quality of sequencing was evaluated with FastQC (v0.11.9)[81] and summarized with MultiQC (v1.13)[82]. Reads were aligned to the mouse reference genome (mm10) in local mode with Bowtie2[83](v2.4.1), using – X 1000 to specify an increased pair distance to 1000bp. Samtools (v1.15.1)[84] and Picard (v2.26.11; http://broadinstitute.github.io/picard/) were used to sort, deduplicate and index reads. Peaks were called with MACS3 (v3.0.0a7; https://github.com/macs3-project/MACS)[85] and specifying --nomodel and --nolambda for the ’callpeaks()’ command. Peaks overlapping mm10 blacklisted/block listed regions called by ENCODE[86,87] and in the original ATAC-seq paper[36] were also removed with BEDTools (v2.30.0)[88].

For visualization with IGV, IGVTools (v2.15.2) was used to convert read pileups to TDFs. The fraction of reads in peaks was calculated with DeepTools (v3.5.1)[89] using the plotEnrichment command. The average mapping distance flag was extracted from the SAM files with a custom script available at our GitHub repo (https://github.com/sarahmcclymont/SN4741_ATAC/) to generate the fragment length plot. Mouse (mm10) transcriptional start site (TSS) coordinates were downloaded from the UCSC Genome Browser[90] (Mouse genome; mm10 assembly; Genes and Gene Predictions; RefSeq Genes track using the table refGene), and DeepTools (v3.5.1)[89] was used to plot the pileup of reads overtop of these TSSs. Conservation under peaks (phastCons)[91] and the genomic distribution of peaks were calculated using the Cis-regulatory Element Annotation System (CEAS)[92] and conservation tool of the Cistrome[93] pipeline. Analysis can be found at http://cistrome.org/ap/u/smcclymont/h/sn4741-atac-seq-ceas-and-conservation.

### ATAC-seq Normalization and Differential Peak Analysis

Each sample’s peak file and BAMs were read into and analysed with DiffBind (v3.8.1)[48]. Peaks present in two or more libraries were considered in the consensus peakset. Reads overlapping these consensus peaks were counted with ’dba.count()’ specifying summits = 100, bRemoveDuplicates = TRUE. These read counts were normalized with ’dba.normalize()’ on the full library size using the RLE normalization method as it is native to the DESeq2 analysis we employed in the following ’dba.analyze()’ step. The volcano plot was generated using a custom script using the output of the ’dba.report()’ command, where th = 1 and fold = 0 and bCounts = T, to output all peaks regardless of their foldchange or significance. Significantly differentially accessible regions (filtered for abs(Fold)>1 & FDR < 0.05) were submitted to GREAT (v4.0.4)[94] and the gene ontology of the nearest gene, as identified with the basal+extension method (where proximal was considered to be ±5kb) was assessed and plotted. **ATAC-seq comparison to *ex vivo* MB and FB DA neurons**

Previously generated ATAC-seq libraries from *ex vivo* E15.5 mouse embryonic DA neurons[50] were re-analyzed in parallel, following all the above alignment and filtering steps. DiffBind (v3.8.1) was used to compare the samples, as above, and the R package UpSetR (v1.4.0)[95] was used to plot the overlap of peaks between conditions.

### Bulk RNA-seq Library Preparation, Sequencing, and Alignment

Cells were run through QIAshredder (Qiagen) and total RNA was extracted using the RNeasy Mini Kit (Qiagen) according to the manufacturer’s recommendations, except that RNA was eluted twice in 50μL of water. Total RNA integrity was determined with the RNA Pico Kit (Agilent) on the Agilent 2100 Bioanalyzer. RNA samples were sent to the Johns Hopkins University Genetic Resources Core Facility (GRCF) for library prep (NEBNext Ultra II directional library prep kit with poly-A selection) and sequencing. The libraries were pooled and sequenced on an Illumina NovaSeq 6000 (SP flow cells), using 2×50 bp reads per library, to a combined depth of 1.6 billion reads. The quality of sequencing was evaluated via FastQC. FASTQ files were aligned to the mouse reference genome (mm10) with HISAT2[96] (v2.0.5) and sample reads from different lanes were merged using samtools[84](v.1.10) function “merge.”. Aligned reads from individual samples were quantified against the mm10 reference transcriptome with the subread[97–99](v1.6.1) function “featureCounts” [100], using -t exon and -g gene_id, (**Supplemental Figure 3A**).

### Bulk RNA-seq Analysis

The DESeq2 (v3.15) package was used for data quality assessment and analyses. A DESeqDataSet of count data was generated using “DESeqDataSetFromMatrix” (design = ∼ temp). The data underwent variance stabilizing transformation (vst) prior to using “plotPCA” to visualize experimental covariates/batch effects (**Supplemental Figure 3B**) and R package “pheatmap” (v1.0.12; https://CRAN.R-project.org/package=pheatmap) to visualize the sample-to-sample distances (**Supplemental Figure 3C**).

Genes with an average of at least 1 read for each sample were analyzed to identify differentially expressed (DE) genes between temperature conditions, using the function “DESeq.” P-value distribution after differential expression (DE) analysis **(Supplemental Figure 3D**) verified that the majority of called DE genes are significant. Results (alpha = 0.01) were generated and subjected to log fold change shrinkage using the function “lfcShrink” (type = “apeglm”)[101] for subsequent visualization and ranking. The function “plotMA” was used to generate MA plots, both before and after LFC shrinkage, to visualize the log_2_ fold changes attributable to the non-permissive temperature shift over the mean of normalized counts for all the samples in the DESeqDataSet (**Supplemental Figure 3E-F**). MA plots demonstrated that log fold change shrinkage of the data successfully diminished the effect size of lowly expressed transcripts with relatively high levels of variability.

Volcano plots were generated using a custom function to visualize log_2_ fold changes of specific genes in the dataset. A gene was considered significantly differentially expressed if it demonstrated an adjusted p-value < 0.01 and |log_2_ FC| > 1.5. These significantly differentially expressed genes were submitted to Enrichr[51–53] for analyses within the “ontologies” and “cell types” categories. The upregulated and downregulated gene sets were passed to STRING [66] for analysis of protein-protein interactions and network relationships.

### Bulk RNA-seq comparison to *ex vivo* MB and FB DA neurons

Read counts from the SN4741 bulk RNA-seq dataset were converted to RPKM and compared to bulk RNA-seq data from previously generated *ex vivo* mouse embryonic DA neurons (NCBI GEO: GSE122450;[50]). A Pearson correlation heatmap was generated using ggplot2[102].

### Promoter capture HiC library generation

PcHiC was performed as previously described[103], with minor modifications. Briefly, SN4741 cells were cultured at the non-permissive temperature and plated at five million cells per 10cm dish. The cells were crosslinked using 1% formaldehyde, snap frozen using liquid nitrogen, and stored at -80°C. The cells were dounce homogenized and restriction enzyme digestion, using 400 units *HindIII-HF* overnight at 37°C. The total volume was maintained at 500µL, through addition of 1X NEBuffer 2.1. Heat inactivation was performed at 80°C for 20 minutes, and biotinylated-dCTP was used for biotin fill-in reaction. Blunt-end ligation was performed using Thermo T4 DNA ligase, with cohesive end units maintained at 15,000 and buffer and water volumes adjusted to ensure a total volume of 665µL was added to each Hi-C tube. Cross-linking was performed overnight, with additional (50μL) proteinase K added for two hours the following day. DNA purification was split across two reactions using 2mL PhaseLock tubes, and volumes were adjusted accordingly. Each PhaseLock reaction was split again into two vials for ethanol purification, and centrifugation at step 6.3.8 was performed at room temperature. The pellets were dissolved in 450µL 1X TLE and transferred to a 0.5mL 30kD Amicon Column. After washing, the column was inverted into a new container, and no additional liquid was added to raise the volume to 100µL. All four reactions were combined, the total volume determined, and RNAseA (1mg/mL) equal to 1% of the total volume was added for 30 minutes at 37°C.

The libraries were assessed for successful blunt-end ligation by a ClaI restriction enzyme digest of PCR products, as previously described[103]. Biotin was removed from un-ligated ends and DNA was sheared to a size of 200-300bp using the Covaris M220 (High setting, 35 cycles of 30s “on” and 90s “off”; vortexing/spinning down samples and changing sonicator water every 5 cycles). Size selection was performed using AMpure XP magnetic beads, as previously described[103] except that all resuspension steps were increased by 5µL, so that 5μL could be used for QC with the High Sensitivity DNA Assay (Agilent) on the Agilent 2100 Bioanalyzer (at three stages: post-sonication, post-0.8x size-selection, and post-1.1x size-selection). The remaining protocol was performed as described. Capture probes (Arbor Biosciences; https://github.com/nbbarrientos/SN4741_pcHiC) were designed against mouse (mm10) RefSeq transcription start sites, filtering out “XM” and “XR” annotated genes. The remaining promoters were intersected with the *in silico* digested *HindIII* mouse genome, to retain all *HindIII* fragments containing a promoter. Potential probes sites were assessed ±330bp of the *HindIII* cut site on either end of the fragment and finalized probe sets were filtered using no repeats and “strict” criteria, as defined by Arbor Biosciences. After generating a uniquely indexed HiC library with complete Illumina adapters, probes targeting promoter containing fragments were hybridized following Arbor Biosciences capture protocol (v4) at 65°C, 1µg DNA, and one round of capture. The library was PCR amplified before sequencing on an Illumina NovaSeq 6000 (SP flow cells), using 2×50 bp reads per library, to a combined depth of 1.6 billion reads.

### Promoter capture HiC data analysis

Raw pcHiC reads for each replicate (n=4) were evaluated for quality via FastQC. FASTQ files were mapped to mm10 using Bowtie2[83] (v.2.4.1) and filtered using HiCUP[104](v. 0.8). The HiCUP pipeline was configured with the following parameters: FASTQ format (Format: Sanger), maximum di-tag length (Longest: 700), minimum di-tag length (shortest: 50), and filtering and alignment statistics were reported (**Supplemental Figure 4A-C**). BAM files were generated for each replicate using samtools[84](v.1.10). DeepTools[89](v.3.5.1) before read coverage similarities and replicate correlation was assessed using the function “multiBamSummary” (in *bins* mode) to analyze the entire genome. A Pearson correlation heatmap was generated using the function “plotCorrelation” (**Supplemental Figure 4D**). As a result of high Pearson correlation coefficient among replicates (r > 0.93), library replicates were combined. The CHiCAGO[105](v. 1.18.0) pipeline was used to convert the merged BAM file into CHiCAGO format. The digested mm10 reference genome was used to generate a restriction map file, a baited restriction map file, and the rest of required input files (.npb, .nbpb, and .poe) required to run the CHiCAGO pipeline.

### AVAILABILITY OF DATA AND MATERIALS

All data and analysis pipelines are available at https://github.com/rachelboyd. ATAC-sequencing, RNA-sequencing, and promoter-capture Hi-C data will be available at the Gene Expression Omnibus (GEO) under the accession number GSEXXXXXX upon manuscript acceptance.

## ABBREVIATIONS

ActB: β-actin
Aldh1a1: Aldehyde Dehydrogenase 1 Family Member A1
ATAC-seq: Assay for Transposase-Accessible Chromatin using Sequencing
BAM: Binary Alignment and Map
Cacna1b: Calcium channel, voltage-dependent, N type, alpha 1B subunit
Ccna2: Cyclin A2
Cdc6: Cell division cycle 6
Cdh13: Cadherin 13
CEAS: Cis-Regulatory Element Annotation System
Celf5: CUGBP Elav-Like Family Member 5
Cenpf: Centromere protein F
Cntn2: Contactin 2
CO2: Carbon Dioxide
CRE: Cis Regulatory Element
DA: Dopaminergic
DMEM: Dulbecco’s Modified Eagle Medium
Dpysl5: Dihydropyrimidinase-like 5
E13.5/15.5: Embryonic Day 13.5/15.5
ENCODE: Encyclopedia of DNA Elements
FB: Forebrain
FBS: Fetal Bovine Serum
FDR: False Discovery Rate
Foxa2: Forkhead Box A2
Gins1: GINS complex subunit 1 (Psf1 homolog)
GO: Gene Ontology
GRCF: Genetics Core Research Facility
GWAS: Genome-Wide Association study
Hdac11: Histone deacetylase 11
Hmga2: High Mobility Group AT-Hook 2
Id2: Inhibitor Of DNA Binding 2
IGV: Integrative Genomics Viewer
Il33: Interleukin 33
Irx3: Iroquois Homeobox 3
KEGG: Kyoto Encyclopedia of Genes and Genomes
LFC: Log Fold-Change
Lmx1b: LIM Homeobox Transcription Factor 1 Beta
MB: Midbrain
Mki67: Marker of Proliferation Ki-67
Nes: Nestin
Nova2: NOVA alternative splicing regulator 2
Nr4a2: Nuclear Receptor Subfamily 4 Group A, Member 2
Nrx1: Neurexin 1
Ntrk1: Neurotrophic receptor tyrosine kinase 1
OCR: Open Chromatin Region
Olig3: Oligodendrocyte transcription factor 3
PC(A): Principal Component (Analysis)
pcHi-C: Promoter-Capture Hi-C
PD: Parkinson Disease
Pitx3: Paired-like homeodomain 3
Ptgds: : Prostaglandin D2 synthase
(q)PCR: (Quantitative) Polymerase Chain Reaction
QC: Quality Control
RNA: Ribonucleic Acid
RPKM: Reads per kilobase of exon per million reads mapped
RT: Reverse Transcriptase
Scn1b: Sodium Voltage-Gated Channel Beta Subunit 1
scRNA-seq: Single Cell RNA sequencing
Slc6a3: Solute Carrier Family 6 Member 3
SN: Substantia Nigra
SNCA/Snca: Alpha-synuclein
SV40Tag: Simian Virus 40 T antigen
TH/Th: Tyrosine Hydroxylase
Tmem179: Transmembrane protein 179
ts: Temperature-Sensitive
TSS: Transcriptional Start Site
Unc13a: Unc-13 homolog A
vst: Variance Stabilizing Transformation

## ACKNOWLEDGEMENTS

The authors would like to acknowledge Ernest Arenas (Karolinska Institutet), for providing SN4741 cells, as well as the Johns Hopkins Genomics Core Research Facility (GCRF) and the WiCell Research Institute, for providing technical services.

## FUNDING

This research, undertaken at Johns Hopkins University School of Medicine, was supported in part by awards from the National Institutes of Health (NS62972 and MH106522) to A.S.M., by T32 GM007814-40 to R.J.B. and N.B.B., and by the Canadian Institutes of Health Research (DFD-181599) to R.J.B.

## AUTHOR INFORMATION

## Contributions

New data was generated by S.A.M., P.W.H., W.D.L., and E.W.L.; analyzed by R.J.B., S.A.M.,

P.W.H., N.B.B., W.D.L., and A.S.M. The manuscript was written by R.J.B., S.A.M., N.B.B., W.D.L., and A.S.M. Figures were created by R.J.B., S.A.M., and N.B.B. All authors reviewed and approved the manuscript.

### ETHICS DECLARATIONS

#### Ethics approval and consent to participate

Not applicable.

#### Consent for publication

Not applicable.

#### Competing interests

The authors declare no competing interests.

## Notes

https://github.com/rachelboyd

### Competing Interest Statement

The authors have declared no competing interest.

### Summary of Updates

Changes to figures/captions and in-text figure references.

