## Supplemental Table 1 and Figures 1-4 for "Evaluating the mouse neural precursor line, SN4741, as a suitable proxy for midbrain dopaminergic neurons"

**Supplemental Table 1:** RT-qPCR primers for testing expression of dopaminergic neuron markers.

| Gene | Forward Primer | Reverse Primer | Expected Amplicon |
|---|---|---|---|
| *Th* | CTGTCCACGTCCCCAAGGTTCA | CAATGGGTTCCCAGGTTCCG | 147 bp |
| *Foxa2* | CCCTACGCCAAATGAACTCG | GTTCTGCCGGTAGAAAGGGA | 221 bp |
| *Nr4a2* | GTGTTCAGGCGCAGTATGG | TGGCAGTAATTTCAGTGTTGGT | 153 bp |
| *Slc6a3* | GAGGCCCGATAAGAGCTCAAG | CCTTCTTCTTCGACTGCCTCC | 111 bp |
| *Actb* | TGGCTCCTAGCACCATGAAG | AGCTCAGTAACAGTCCGCCTA | 188 bp |

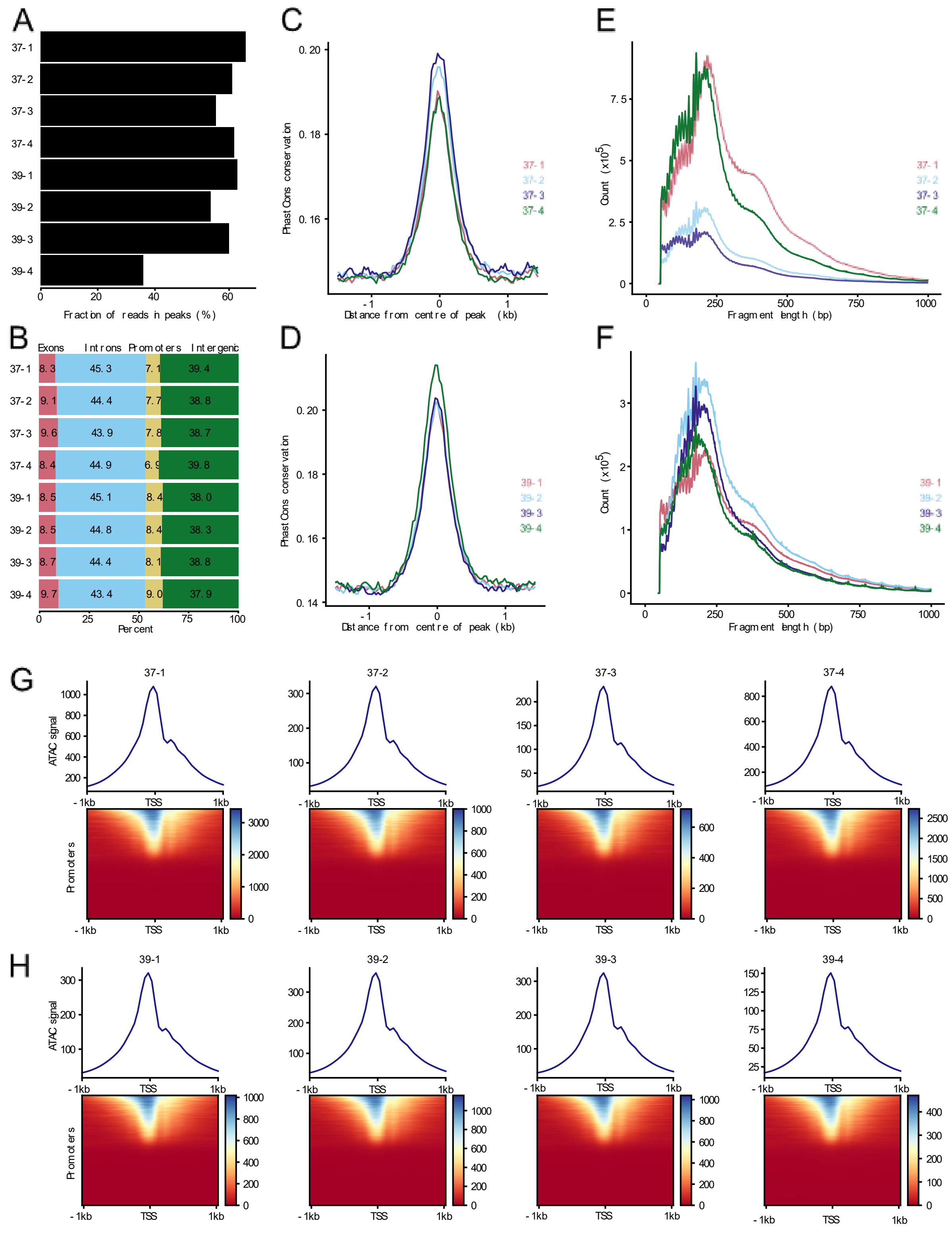


**Supplemental Figure 1: ATAC-seq libraries are technically and biologically relevant. A**) The ATAC-seq libraries have low levels of background reads, with most reads falling in peaks. **B**) Peaks are distributed evenly between all samples, with the majority of peaks falling over non-coding regions of DNA (intergenic and intronic). **C, D**) Sequences underlying peaks are well conserved, suggesting evolutionary constraint of these sequences (i.e.: a suggestion of biological functionality). **E, F**) Libraries all demonstrate the characteristic nucleosome patterning and the 10bp DNA pitch sawtooth pattern, indicating proper transposition. **G, H**) Libraries all have an enrichment of reads over transcriptional start sites (TSS).

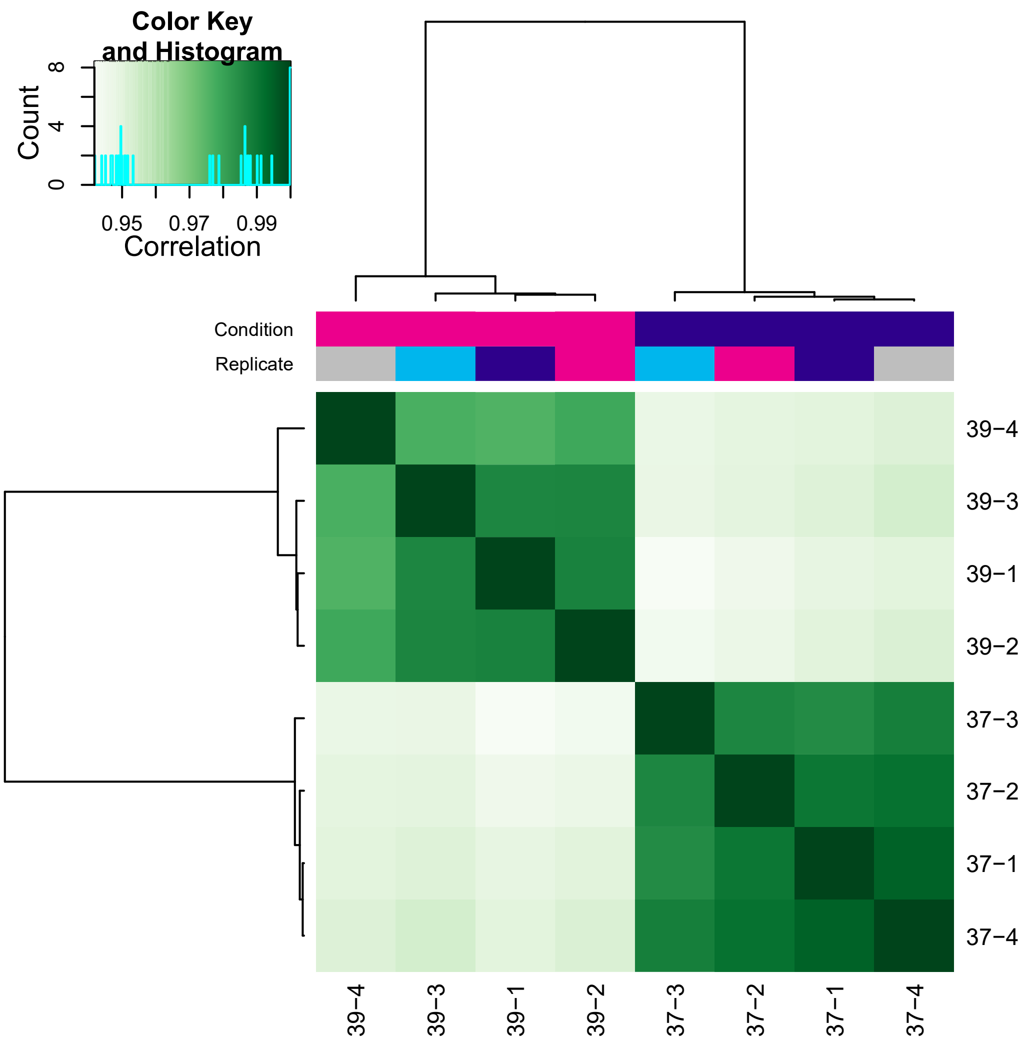


**Supplemental Figure 2: ATAC-seq libraries are well correlated within and between temperature conditions.** The ATAC-seq libraries are highly similar between replicates ($r^2 > 0.97$). The libraries are less, but still highly, correlated between temperature conditions ($0.94 < r^2 > 0.96$).

A

| | 37-1 | 37-2 | 37-3 | 37-4 | 39-1 | 39-2 | 39-3 | 39-4 |
|---|---|---|---|---|---|---|---|---|
| Total alignments | 97,619,380 | 77,429,158 | 185,166,944 | 115,299,901 | 113,843,949 | 220,871,735 | 188,079,600 | 132,067,167 |
| Assigned | 73,236,573 | 57,769,924 | 137,826,794 | 86,026,485 | 86,612,965 | 167,241,485 | 142,495,717 | 99,803,070 |
| Unassigned_MultiMapping | 16,954,447 | 13,492,419 | 32,626,596 | 20,033,741 | 17,064,810 | 33,829,184 | 28,357,700 | 20,172,825 |
| Unassigned_NoFeatures | 2,783,817 | 2,408,417 | 5,842,993 | 3,585,468 | 4,302,522 | 8,348,391 | 7,367,554 | 5,008,989 |
| Unassigned_Ambiguity | 2,391,543 | 1,878,742 | 4,464,367 | 2,773,548 | 2,963,036 | 5,776,109 | 4,786,564 | 3,511,402 |
| Unassigned_Unmapped | 2,253,000 | 1,879,656 | 4,406,194 | 2,880,659 | 2,900,616 | 5,676,566 | 5,072,065 | 3,570,881 |
| Unassigned_MappingQuality | 0 | 0 | 0 | 0 | 0 | 0 | 0 | 0 |
| Unassigned_Chimera | 0 | 0 | 0 | 0 | 0 | 0 | 0 | 0 |
| Unassigned_FragmentLength | 0 | 0 | 0 | 0 | 0 | 0 | 0 | 0 |
| Unassigned_Duplicate | 0 | 0 | 0 | 0 | 0 | 0 | 0 | 0 |
| Unassigned_Secondary | 0 | 0 | 0 | 0 | 0 | 0 | 0 | 0 |
| Unassigned_Nonjunction | 0 | 0 | 0 | 0 | 0 | 0 | 0 | 0 |
| Unassigned_Overlapping_Length | 0 | 0 | 0 | 0 | 0 | 0 | 0 | 0 |

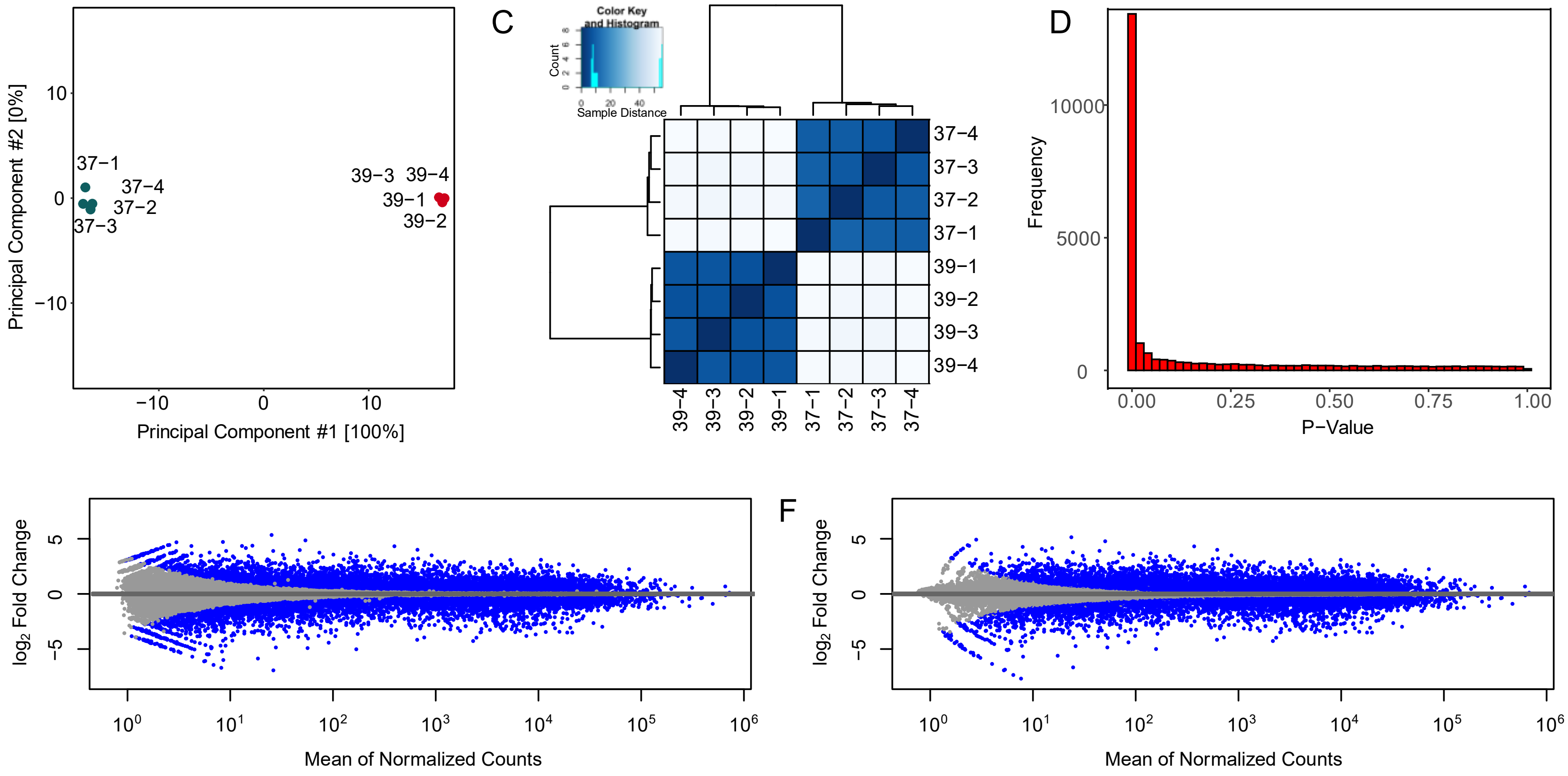


**Supplemental Figure 3. Bulk RNA-seq Analysis: A)** Summary of counting results using featureCounts, including the number of successful alignments assigned and number of alignments that could not be assigned due to various filters to the reference annotation. **B)** PCA plot of the four temperature condition replicates (blue, 37°C; 39°C, red). **C)** Heatmap and hierarchical clustering based on sample-sample distances. **D)** P-value distribution after differential gene expression analysis. **E-F)** MA-plots of log2 fold change versus mean of normalized counts with DESeq2 before (E) and after (F) Log fold change shrinkage. Blue = genes/transcripts that are statistically differentially expressed (P-Value < 4e-7).

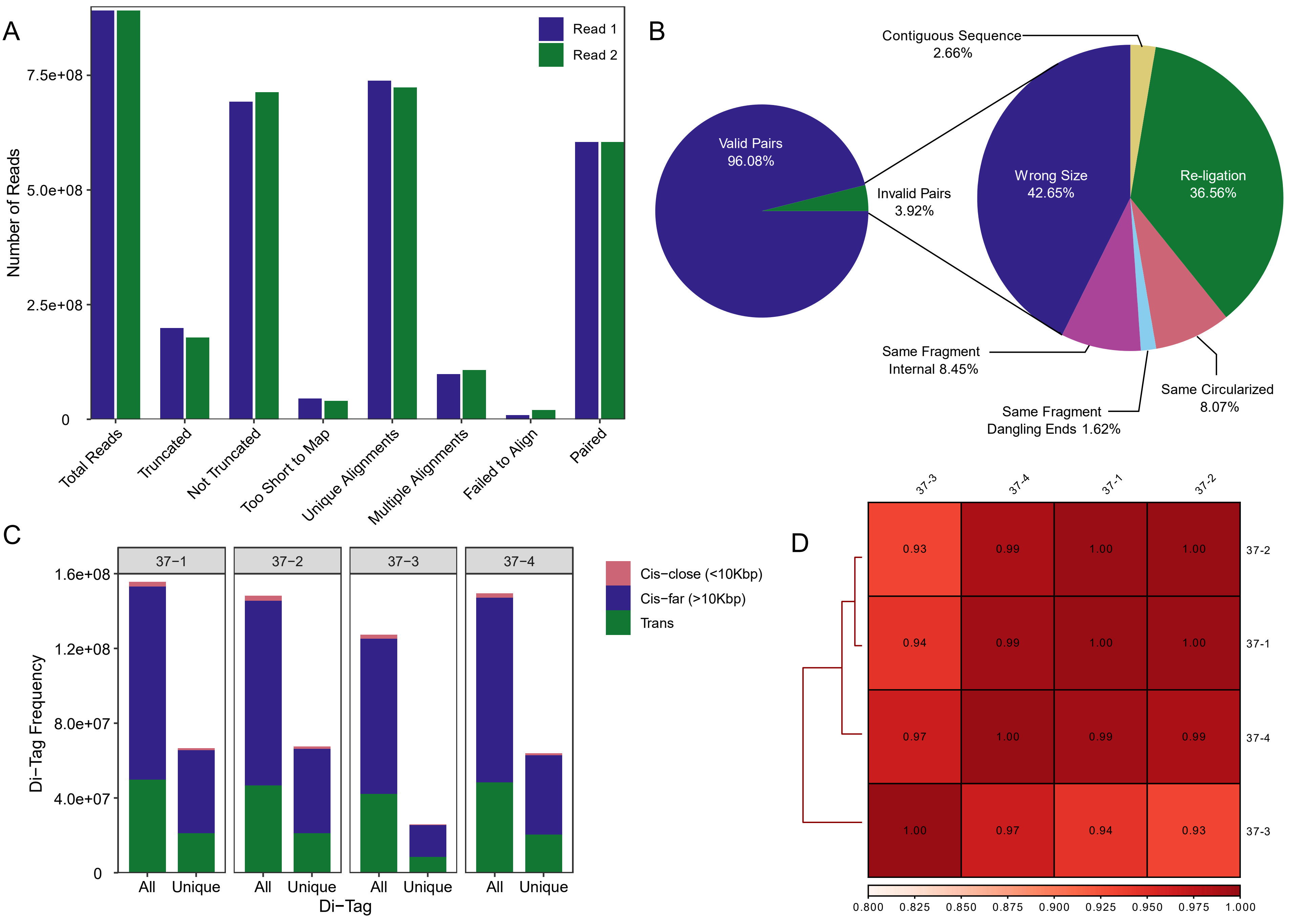

A
Number of Reads
Read 1
Read 2
7.5e+08
5.0e+08
2.5e+08
0
Total Reads
Truncated
Not Truncated
Too Short to Map
Unique Alignments
Multiple Alignments
Failed to Align
Paired
B
Valid Pairs
96.08%
Invalid Pairs
3.92%
Contiguous Sequence
2.66%
Wrong Size
42.65%
Re-ligation
36.56%
Same Fragment
Internal 8.45%
Same Fragment
Dangling Ends 1.62%
Same Circularized
8.07%
C
Di−Tag Frequency
37−1
37−2
37−3
37−4
1.6e+08
1.2e+08
8.0e+07
4.0e+07
0
All
Unique
Di−Tag
Cis−close (<10Kbp)
Cis−far (>10Kbp)
Trans
D
37-3
37-4
37-1
37-2
0.93
0.99
1.00
1.00
0.94
0.99
1.00
1.00
0.97
1.00
0.99
0.99
1.00
0.97
0.94
0.93
0.800
0.825
0.850
0.875
0.900
0.925
0.950
0.975
1.000

**Supplemental Figure 4.** ***in silico*** **quality control metrics for all Promoter-capture Hi-C libraries with HiCUP**: **A)** Bar chart showing the number of reads (for all replicates combined) that required truncation prior to alignment and the number of reads that were uniquely aligned, too short to map, had multiple alignments, failed to align, and were paired reads. **B)** Pie chart showing di-tag distribution of valid versus invalid fragments after filtering of Hi-C artifacts. Invalid di-tags are those that originated from (1) circularized (self-ligated) fragments, (2) fragments with only one end overlapping a restriction cut site (dangling ends), (3) fragments with neither end overlapping a restriction cut site (internal), (4) re-ligated fragments, (5) fragments made of contiguous sequences, or (6) di-tags of the wrong size. Valid pairs that passed filtering accounted for 96.1% of di-tags captured. **C)** Bar chart showing the number of unique di-tags after de-duplication. During PCR amplification a single di-tag can be amplified multiple times giving rise to multiple duplicates of the same fragment. HiCUP removes all but one copy of these duplicates for subsequent analyses. **D)** Correlation heatmap showing the Pearson Correlation coefficient among four pcHiC library replicates for SN4741 cells at the non-permissive temperature of 37°C.
